# Investigating Antimicrobial Resistance Genes in Kenya, Uganda and Tanzania Cattle Using Metagenomics

**DOI:** 10.1101/2023.11.07.565943

**Authors:** Kauthar M. Omar, George L. Kitundu, Adijat O. Jimoh, Dorcus N. Namikelwa, Felix M. Lisso, Abiola A. Babajide, Seun E. Olufemi, Olaitan I. Awe

## Abstract

Antimicrobial resistance (AMR) is a growing problem in African cattle production systems, posing a threat to human and animal health and the associated economic value chain. However, there is a poor understanding of the resistomes in small-holder cattle breeds in East African countries. This study aims to examine the distribution of Antimicrobial Resistance Genes (ARGs) in Kenya, Tanzania, and Uganda cattle using a metagenomics approach.

We used the SqueezeMeta-Abricate (assembly-based) pipeline to detect ARGs and benchmarked this approach using the Centifuge-AMRplusplus (read-based) pipeline to evaluate its efficiency. Our findings reveal a significant number of ARGs of critical medical and economic importance in all three countries, including resistance to drugs of last resort such as carbapenems, suggesting the presence of highly virulent and antibiotic-resistant bacterial pathogens (ESKAPE) circulating in East Africa.

Shared ARGs such as aph(6)-id (aminoglycoside phosphotransferase), tet (tetracycline resistance gene), sul2 (sulfonamide resistance gene) and cfxA_gen (betalactamase gene) were detected. Assembly-based methods revealed fewer ARGs compared to read-based methods, indicating the sensitivity and specificity of read-based methods in resistome characterization.

Our findings call for further surveillance to estimate the intensity of the antibiotic resistance problem and wider resistome classification. Effective management of livestock and antibiotic consumption is crucial in minimizing antimicrobial resistance and maximizing productivity, making these findings relevant to stakeholders, agriculturists, and veterinarians in East Africa and Africa at large.

## Introduction

According to WHO, antimicrobials are used for the prevention and/or treatment of infections in animals, humans, and plants. Antimicrobials include antibiotics, antifungals, and antiparasitics (WHO, 2021). Antimicrobial resistance (AMR) refers to the ability of microorganisms such as bacteria, fungi, parasites and viruses to evolve and become resistant to antibiotics, and when this happens, infections become more difficult to treat, thereby increasing the risk of disease spread, severe illness, and ultimately death (Prestinaci *et al*., 2015). This has become a global health concern worldwide (Donado-Godoy *et al*., 2015). The emergence of resistance to antibiotics in microorganisms, especially bacteria, is partially due to the existence of resistance genes (Zalewska *et al*., 2021), and farm animals, such as cattle being the prime reservoirs for these genes (Kim and Ahn, 2022).

Despite the importance of antimicrobials in the livestock sector, with reference to veterinary medicines used for the treatment of animal infections and growth promotion of animals, the uncontrolled, subtherapeutic doses of antibiotics in cattle farms can promote the growth of bacteria that are antibiotic resistant, which in turn affects the changes in microbial metabolism of the animal’s rumen and the global health crisis associated with antibiotic resistance (Maron *et al*., 2013; Economou and Gousia, 2015). The transmission of antibiotic resistance genes (ARGs) occurs majorly through conjugation, which refers to horizontal gene transfer that exchanges mobile genetic elements, such as plasmids and transposons that code for ARGs between bacterial species (Redondo-Salvo *et al*., 2020). Bacteria can also develop antibiotic resistance through chromosomal mutation that empowers them with the ability to withstand high concentrations of antimicrobials from recurrent exposure (Baym *et al*., 2016). In other cases, they have natural resistance due to the presence of an impermeable cell membrane or lack of an inherent target molecule for the antibiotic to counter against (Mann *et al*., 2021).

In Africa, reports indicate that the continent has the lowest antimicrobial usage in animals worldwide (WOAH, 2020), even though a high incidence of resistance to antimicrobials have been observed in foodborne pathogens isolated from animals and animal products (Van Boeckel *et al*., 2019). However, there is limited data in East Africa on antimicrobial knowledge, attitudes, and practices (KAP) (Kemp *et al*., 2021). It has been recorded that the extensive usage of antimicrobials/antibiotics for treatment of dairy cattle infected with certain pathogens encourages the development of resistant strains in such host animals (Vincent *et al*., 2018). Research shows that in East Africa, AMR is linked to human-animal contact, community transmission of resistant bacteria and ease of access to cheap antibiotics (Omulo *et al*., 2015) but there is little information on the source, diversity, as well as the distribution of antimicrobial resistant genes in the most non-cultivable environmental bacterial-pathogens in Africa.

The shown ability to project AMR from genetic sequencing data is a significant leap in resistome surveillance. There have been efforts to model the evolution of viral pathogens like SARS-CoV-2 using genomic sequence data (Awe *et al*., 2023), HIV-1 evolution in sub-Saharan Africa (Obura *et al*., 2022), biomarker discovery (Chikwambi *et al*., 2023; Nyamari *et al*., 2023; El Abed *et al*., 2023), malaria/CoVID-19 biomarker discovery (Nzungize *et al*., 2022), analysis of RNA-seq and ChIP-seq data (Ather et al., 2018), Ebola Virus comparative genomics (Oluwagbemi and Awe, 2018). Genomics and bioinformatics have been reported to be playing a key role in Newborn screening (Wesonga and Awe, 2022) and in agriculture. For instance, there have been computational methods to define gene families and their expression in legumes (Die *et al*., 2019). High sensitivity, which is the ability to recognize AMR determinants associated with antimicrobial resistance phenotype and high specificity, which is the ability to recognize the absence of AMR determinants in an antimicrobial susceptible phenotype have been observed depending on the bacterial species studied (Hendriksen *et al*., 2019). One of the tools that can be used in analysing sequencing DNA sequences is metagenomics. Metagenomics refers to the study of the structure and function of the total nucleotide sequences which is isolated and analysed from the entire community of micro-organisms present in a sample (Thomas *et al*., 2012). A notable technique called shotgun metagenomics, studies the genetic content of an environmental sample by extracting and sequencing its microbial community’s total genomic DNA. As a result, this technique yields a substantial overview of the microbiota and allows for the simultaneous investigation of the taxonomic classification and functional aspects of the microbial communities (Kibegwa *et al*., 2020), as compared to its counterpart, amplicon sequencing approach which only targets a particular region of the community’s genomic DNA. Hence, shotgun metagenomics is the most suitable approach for our study.

According to the East Africa Integration agenda, the East African Community (EAC) aims at transforming the region into a single market (Umulisa, 2020). This means goods and services will be moving across different partner states and that includes the most widely used dairy products from cattle. For milk alone, Eastern Africa is recorded as the first and leading milk-producing region in Africa. It represents 68% of the continent’s milk production with Kenya and Tanzania being amongst the biggest dairy producers in Africa (Bingi and Tondel, 2015). All of this, when integrated into the value chain, creates employment opportunities through businesses done by the movement of products around the member states.

An understanding of circulating resistomes is thus crucial since cattle management becomes quite difficult with disease burden, leading to a reduced production as well as increasing production costs, which threatens the economic growth agenda of the EAC. In this case, it is important to understand the emergence of resistomes within the states implicated to better manage livestock in maximizing production and lowering risk and production costs. In addition to the cost-related benefits of the study, the outcomes will assist stakeholders in a large extent to improve the standardization protocols that control the circulation of cattle-derived products and guide veterinarians on informed decisions on how to efficiently manage clinical cases.

## Methods

### Workflow

#### Data Acquisition

Metagenomics data of Kenyan, Ugandan and Tanzanian cattle gut microbiome was obtained from public repositories. The data was obtained using the search method described in Supplementary methods: Data Search. The Kenyan and Ugandan datasets were retrieved from ENA under bioprojects PRJEB28482 and PRJEB20456 respectively, using fasterq-dump (ncbi, n.d.). All samples from bioprojects PRJEB28482 (Kenya dataset) were used but only six samples from the bioproject PRJEB20456 (Uganda dataset) were used, samples ERR1950666 to ERR1950671.This was because Uganda dataset had a significantly higher read depth per sample than Kenya and Tanzania datasets. The grabseqs tool (Taylor *et al*., 2020) was used to retrieve the Tanzania dataset from MG-RAST server (Meyer *et al*., 2008) under the project mgp81260. Table 1 highlights the total number of samples and reads obtained from each country.

**Table 1:**
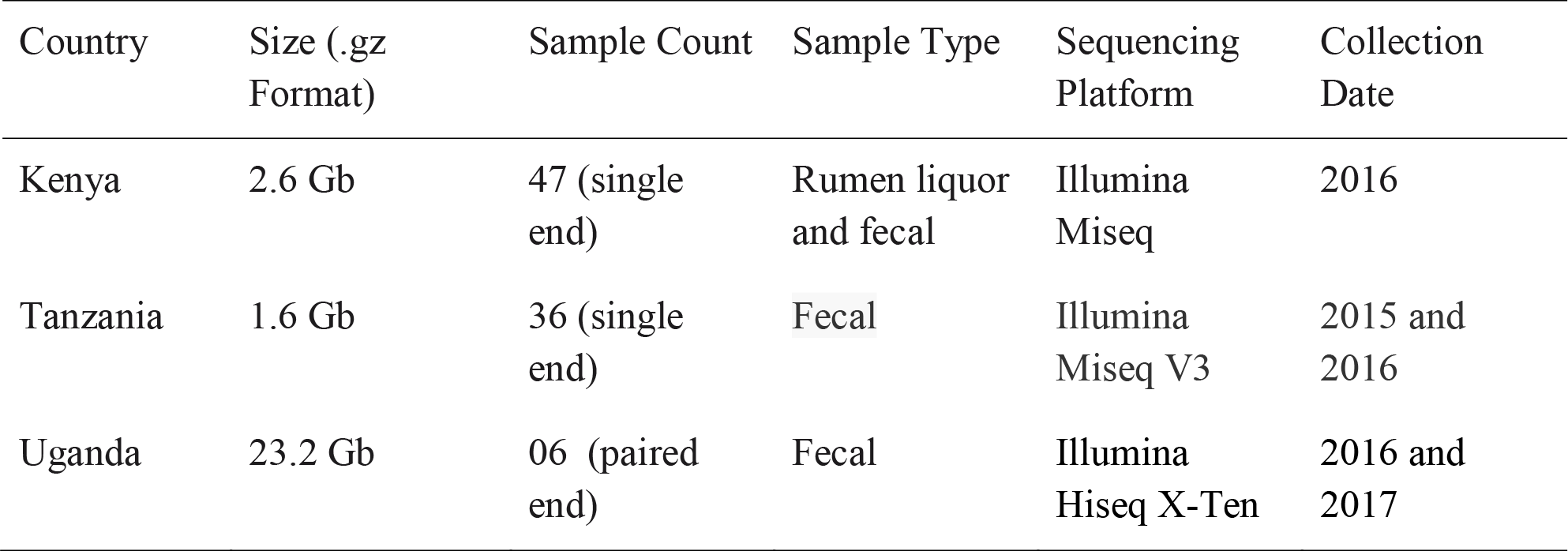
Metadata describing selected experimental aspects of the samples obtained from each country.

### Quality Control

Fastqc was used to assess the quality of the raw reads (Andrews, 2010). Multiqc was used to generate an aggregate report for all fastqc reports per country (Ewels *et al*., 2016). The retrieved reads were all of good quality i.e a Q>20, thus, we saw no need to further trim them to much higher qualities since from Q>=20 is within tolerable limit for Illumina data without interfering with subsequent downstream analyses.

### Metagenomics Data Analysis

Metagenomics analysis of taxonomic abundance and functional annotation was achieved with SqueezeMeta v1.5.1, Jan 2022 (Tamames & Puente-Sánchez, 2019), which is a pipeline that constitutes all the steps from assembly of reads to functional annotation. Megahit was used for assembling (Li *et al*., 2015). Prinseq was used to remove short contigs (<200 bps) (Schmieder and Edwards, 2011). Contig statistics were determined with prinseq (Schmieder and Edwards, 2011). Barrnap was used to predict RNAs (Seemann, 2014). RDP classifier was used to taxonomically classify 16S rRNA sequences (Wang *et al*., 2007). Aragorn was used for predicting tRNA/tmRNA sequences (Laslett, 2004). Prodigal was used to predict open reading frames (ORFs) (Hyatt *et al*., 2010). Diamond (Buchfink *et al*., 2014) was used to search for similarity between eggNOG (Huerta-Cepas *et al*., 2015), GenBank (Clark *et al*., 2015), and KEGG (Kanehisa, 2000). Searches for HMM homology were performed using HMMER3 (Eddy, 2009) for the Pfam database (Finn *et al*., 2015). Using Bowtie2 (Finn *et al*., 2015), reads were mapped against contigs. Binning was conducted using MaxBin2 and Concoct (Wu *et al*., 2015), and Metabat2 (Kang *et al*., 2019).

### Detection of Antimicrobial Resistance Genes (ARGs)

ARGs were detected from the assembly output of the SqueezeMeta pipeline with abricate (tseemann, 2020) using all built in databases which include CARD (Comprehensive Antibiotic Resistance Database) and resfinder, NCBI AMRFinderPlus, ARG-ANNOT, EcOH, MEGARES 2.00 (Feldgarden *et al*., 2019; McArthur *et al*., 2013; Bortolaia *et al*., 2020; Gupta *et al*., 2014; Donado-Godoy et al., 2015; Ingle et al., 2017), which is the default setting for robustness and sensitivity to ARGs detection.

### Benchmarking

To benchmark our analysis workflow (Fig 1a) containing SqueezeMeta for metagenomics analysis and abricate for ARG’s identification, we compared the efficiency of our workflow (Fig 1a) with our benchmarking workflow (Fig 1b). The benchmarking workflow contained AMRplusplus (Gray and Head, 2008) for resistome identification, centrifuge (Kim *et al*., 2016) for taxonomic assignment and Krona (Ondov *et al*., 2011) for visualisation of taxonomic profiles. It is worth mentioning that generally (Fig 1a) workflow was assembly based and (Fig 1b) workflow is read based. In this way we were also assessing, at least in part of our study, the efficiency of assembly-based methods against read based methods in identification of ARGs in our study.

**Fig 1:**
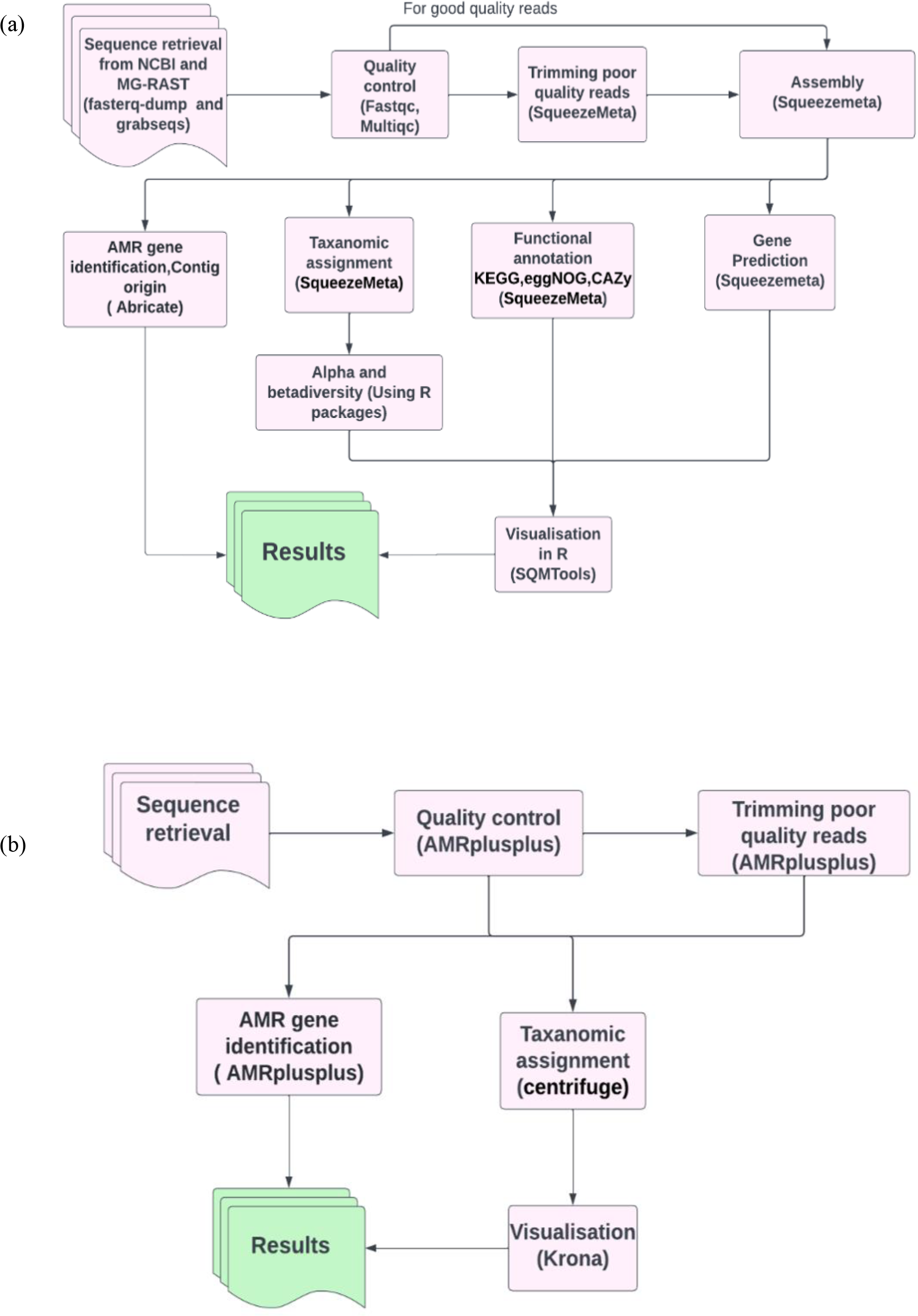
Pictorial representation of the workflows used. (a) Analysis workflow. The workflow begins with sequence retrieval using fasterqdump for Kenya and Uganda dataset and grabseq for the Tanzanian dataset. Quality control, trimming and metagenomic analysis i.e., assembly, taxonomic assignment and functional annotation was carried out in the Squeezemeta pipeline. Visualization of taxonomic assignment and functional annotation was done with SQMTools,an R package in R. AMR gene identification from the assembled contigs was done with Abricate. (b) Benchmarking workflow. In the benchmarking pipeline quality assessment and control, trimming and AMR gene identification was done using tools in the AMRplusplus pipeline. AMRplusplus mines AMR genes directly from the reads. Taxonomic assignment of the reads was done separately using centrifuge. The results from centrifuge were visualized using krona.

### Data Visualization

A key stage in data analysis is the interpretation of the obtained results from different sequence analysis tools. This can be done by using plots, graphs, tables etc, depending on the results you have and the question you are trying to answer. For the sake of our study whereby we wanted to present taxonomic and functional annotations and the visualization of the identified drug classes and ARGs identified, we relied heavily on the use of R and Python programming languages as well as Krona for the visualisation of centrifuge in particular

## Results

### Taxonomic Microbial Abundance and Functional Annotation

Assessment of taxonomic composition is essential in any metagenomics study. It provides you with the insights needed to draw necessary conclusion with regard to your study. In our case where we were interested in detecting antimicrobial resistance, annotating the potential taxonomies found in our samples aids in giving an idea of the severity of resistance and possible mechanism of transmission of these genes across different bacteria if at all certain strains linked with horizontal gene transfers are to be identified. In this section, we provide a brief highlight of the taxonomic and functional abundance of our analysis samples.

#### I. Taxonomic Microbial Abundance in Kenya, Tanzania and Uganda

##### Kenya

Our results highlight some of the details regarding the possible taxonomic composition and functional capacity of microbial communities in samples from each country. In Supplementary Fig 1a, a greater proportion of reads were unmapped in the Kenyan dataset. The phyla *Bacteroidetes, Proteobacteria* and *Firmicutes* were observed to be the most abundant in decreasing order, with *Proteobacteria* being most abundant in some samples. All viruses were unclassified here. Similarly, at the family level of prokaryotic abundance, there was a large percentage of unmapped reads as shown in Supplementary Fig. 2a, produced by SqueezeMeta in the co-assembly mode and the most abundant families were *Prevotellaceae* and *Bacteriodaceae. Pseudomonadaceae* was prevalent in a few of the sample reads with a spiked abundance level.

Fig 2a was generated using the custom pipeline (Fig 1b) to benchmark taxonomic assignment of the metagenomic results obtained from SqueezeMeta. At the phylum level, *Proteobacteria* was generally observed to be most abundant as seen in some of the samples from the SquuezeMeta pipeline. However, at the family level, Pseudomonadaceae was observed to be most abundant, which is consistent to what was observed in some samples of the analysis with SqueezeMeta. Here, a better species resolution is observed.

**Fig 2.**
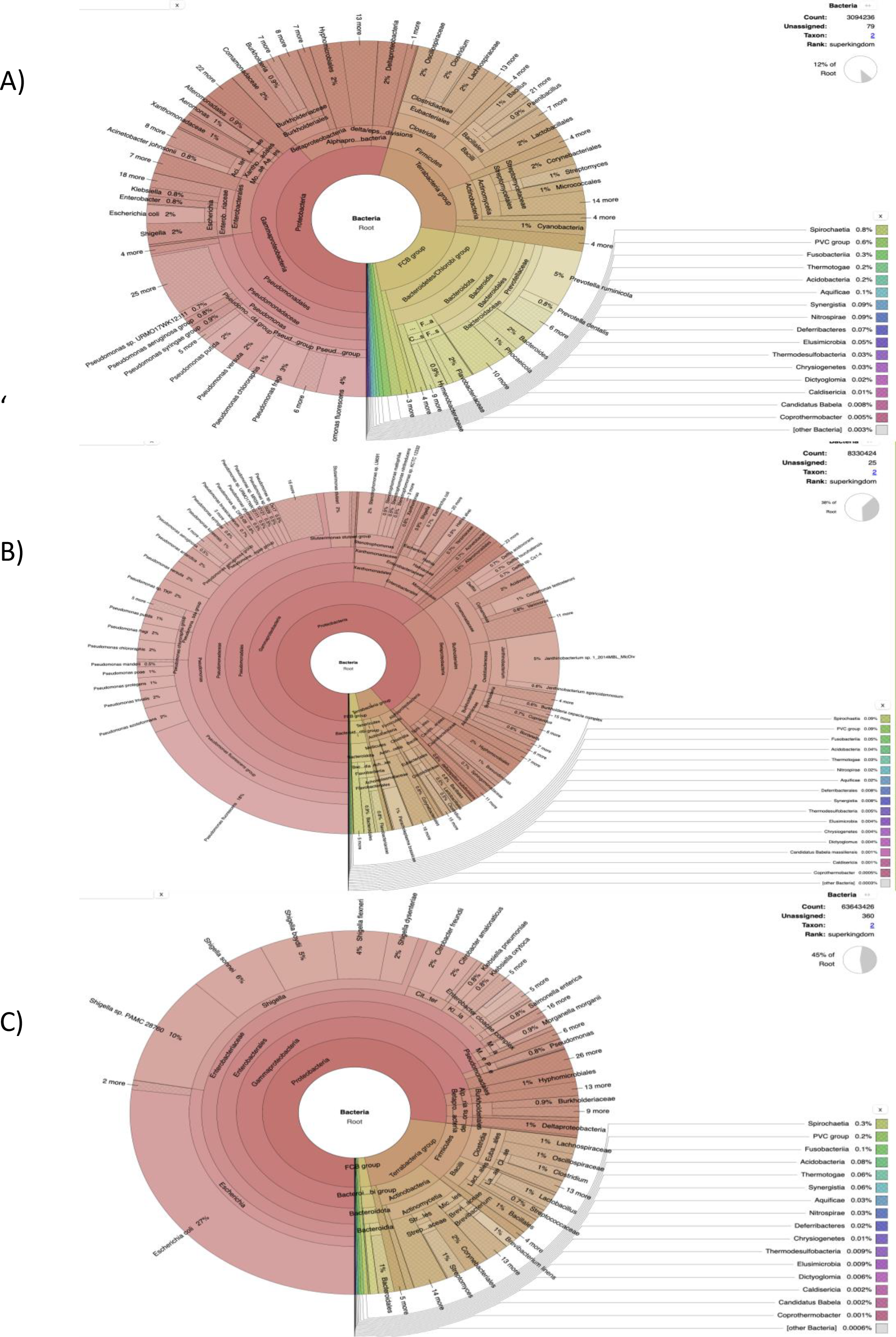
Pie charts describing the taxonomical abundance profile of the cattle rumen liquor and faecal excreta metagenome from Kenya, Tanzania and Uganda for reads that mapped to the centrifuge’s custom bacterial index database. Taxonomical assignment was done directly from the metagenomic reads using centrifuge and the taxonomical classification results were imported into Krona for visualization. The hierarchical taxonomic levels are from Kingdom to species from center to the edges respectively but some phyla are within clade groups like Firmicutes phylum in the Tetrabacteria group (clade). (a) Kenyan samples. (b) Tanzanian samples. (c) Ugandan samples.

##### Tanzania

In the Tanzanian dataset, *Proteobacteria, Firmicutes* and *Bacteroidetes* are the most abundant phyla (Supplementary Fig 1b), a similar observation to that of Kenya. Based on the co-assembly generated by SqueezeMeta, majority of the reads were abundant for the family - *Pseudomonadaceae* in a few samples, followed closely by *Comamonadaceae* and *Oxalobacteraceae* (Supplementary Fig 2b). As seen previously with the Kenyan samples, a high proportion of reads were also unmapped. There were unclassified abundances of bacteria and viruses observed in the taxonomy as well.

The benchmark with Centrifuge i.e., a tool used to implement the custom pipeline, was used to assign taxonomic labels to sequence reads from Tanzania (Fig 2b). The plot also shows species-level assignment as compared to the plot generated from SqueezeMeta. Interestingly, in both plots (Supplementary Fig 1a and Fig 2b), there is a high percentage of the phylum - *Proteobacteria* and unmapped reads as well as a minute level of virus abundance. However, in this plot (Fig 2b), it details and expands more on the *Proteobacteria* phylum, such as the divisions - *Alphaprotobacteria, Gammaproteobacteria*, and the gram-negative bacteria - *Pseudomonas fluorescens, P. azotoformans, Enterobacteriaceae* etc.

#### Uganda

With the Ugandan dataset (Supplementary Fig 1c, there was an uneven distribution of abundances for each phylum in the few samples retrieved from the dataset. The most abundant were *Firmicutes, Proteobacteria* and unmapped or unidentified sequence reads. *Fusobacteria* and *Bacteroides* were only prevalent in a few of them. The most prevalent families observed were *Enterobacteriaceae* and *Oscillospiraceae*.

The benchmark results clearly resolve the most abundant phylum to be *Proteobacteria* (Supplementary Fig 1c and 2c). The taxonomic classification in the figure 2c is similar at the family level with SqueezeMeta results having the most abundant family as *Enterobacteriaceae*. At species level (Fig 2c), there is a presence of ESKAPE pathogens: *Enterobacter spp, Klebsiella pneumoniae* and *Pseudomonas aeruginosa*.

#### II. Functional Annotation Kenya, Tanzania and Uganda Samples

Functional annotation provides an overview of the possible physiological functions that the microbiome might have while they are in the gut environment. In our case this information may be used to understand the possible mechanisms that underlie development of antimicrobial resistance if any. In our study, a total of 15 KEGG orthologue resistance genes were functionally annotated in the Kenyan samples (Supplementary Fig 3a). The analyses show that the orthologue - Iron complex outer membrane receptor protein (K02014) was enriched in a few of the samples, with expression profiles of TPM (Transcripts per million) above 3000, while other genes expressed were below 3000. A similar trend was observed in the Tanzanian samples (Supplementary Fig 3b), with K02014 having high expression levels. On the other hand, there was a disparity of orthologue resistance genes in the Ugandan samples (Supplementary Fig 3c), as there was an absence of K02014. However, there were high gene expression profiles in K06147 (ATP-binding cassette, subfamily B bacterial), with TPM levels above 3000, K07497 (putative transposase) with TPM levels between 2000 and 3000 in a few of the samples. All other gene expression profiles were below 2000.

#### III. Identification of AMR Drug Classes

Understanding the AMR Drug Classes to which potential AMR genes belong to, helps to provide an overview of the extent of resistance to antimicrobials and the potential threat it poses to public health. In this study, four drug classes of resistance common to Kenya, Uganda and Tanzania were identified from the metagenomic assembled contigs. These were Tetracycline, Beta-Lactam, Sulfonamide and Streptomycin. However, there was a varying prevalence to these drug classes in each of the East African countries. *Tetracycline* resistance was the highest in Kenya and Uganda, with of 27% and 24% respectively (Supplementary Figs 4a and 4c). Lincosamide resistance was only detected in the Kenyan cattle samples while Streptomycin, Sulfonamide and Spectinomycin resistance was the lowest at 9% (Supplementary Fig 4A). Carbapenem resistance was found solely in Tanzanian cattle samples. Additionally in Tanzania, resistance levels were lowest to Chloramphenicol and Trimethoprim (both at 6%). Ugandan samples had 15 unique drug classes, including Cephalosporin, Streptothricin, Quinolone, Nitroimadazole and others (Supplementary Fig 4c). The highest number of resistance drug classes was found in Uganda, with a total of 15 (Fig 3a and Table 2).

**Fig 3:** Venn diagram showing the total number of drug classes identified per country uniquely and in comparison, across all three countries from AMR genes mined from assembled contigs. The Venn diagram was generated with bioinformatics and evolutionary genomics Venn diagram tool from Abricate’s output. A total of 24 drug classes were identified in the three countries having 22 in circulation in Uganda, 7 in Tanzania and 6 in Kenya.

**Table 2: Resistance drug classes which were identified in Kenya, Uganda and Tanzania from ARGs mined from assembled contigs.** The table was generated by bioinformatics and evolutionary genomics Venn diagram tool using results from Abricate’s output. This table complements Fig 3a to provide names to the number of drug classes described in the Venn diagram.

On benchmarking our assembly-based approach of ARGs identification with the read-based method in AMRplusplus., we identified a broader spectrum of drug classes. Since the drug classes belonged to ARGs that were directly mined from the reads rather than the assembly (which does not assemble all reads thus leaving out some information). The number of drug classes identified from reads in the 3 countries were reported to be 59. Uganda has all 59 drug classes with 11 unique to Uganda. Tanzania and Kenya had 44 drug classes each. (Supplementary Fig 5 and Supplementary Table 1).

#### IV: Identification of AMR Genes

AMR genes in the microbiome pose a possible threat of unmanageable bacterial infections when the bacteria have virulence factors or acquires virulence factors. In this study, we evaluate the possibility of antibiotic resistance in the case of an infection by identifying AMR genes. In the Ugandan dataset had the highest number of AMR genes detected from the assembly, was 111 (Fig 4 and Table 3). The common AMR genes identified in the three countries that we studied are *aph(6)-id* (aminoglycoside phosphotransferase), *tet* (tetracycline resistance gene), *sul2* (sulfonamide resistance gene) and *cfxA_gen (*betalactamase gene). The two AMR genes unique to Kenya include *cfxA6 (*class A beta lactamase) and *aadA27* (streptomycin/spectinomycin resistance gene) (Table 3).

**Fig 4:** A Venn diagram describing the total number of AMR genes in Kenya, Uganda and Tanzania uniquely as well as comparisons of genes available across the three countries mined from assembled contigs. The Venn diagram was produced using the bioinformatics and evolutionary genomics Venn diagram tool from Abricate’s output. In total, 120 AMR genes were identified in the three countries, with 111 in Uganda, 16 in Tanzania, and 9 in Kenya.

**Table 3: AMR genes present in Kenya, Uganda and Tanzania mined from assembled contigs. counts in individual countries indicate the genes that are unique to that country.** The table was created using the bioinformatics and evolutionary genomics Venn diagram tool under text results from Abricate’s output. This table supplements Fig 4 by giving names to the number of AMR genes that are described in the Venn diagram.

A total of 1425 AMR genes were identified from the reads as a benchmark of the assembly-based approach. Uganda had a total of 1420 ARGs in circulation, 965 of which were unique to Uganda, while Tanzania had 403 AMR genes in circulation with 3 unique to itself. Kenya had 353 genes in circulation which were also found in either Uganda or Tanzania (Supplementary Fig 6 and Supplementary Table 2). There were many genes identified from reads as compared to those mined from assembled contigs. This is because during assembly, assembled contigs exclude some reads which may contain some of the AMR genes.

## Discussion

In our study, we performed metagenomic analysis on public datasets generated from cattle rumen and faecal samples in the three East African countries; Kenya, Tanzania and Uganda. Our data analysis suggests that *Prevotellaceae, Pseudomonadaceae* and *Enterobacteriaceae* are the most abundant families in Kenya, Tanzania and Uganda respectively. These findings agree with previous reports (Kibegwa *et al*., 2020; Okubo *et al*., 2018), also had similar findings. *Enterobacteriaceae*, the most abundant family specifically *Escherichia coli*, the most abundant species in Uganda, *have been* associated with acquiring antimicrobial-resistance-genes from other bacteria through horizontal gene transfer, hence acting as reservoirs of AMR genes (Bailey *et al*., 2010; Carattoli, 2008). In addition to *E. coli*, a higher number of ESKAPE pathogens such as *Enterobacter spp*, Klebsiella *pneumoniae* and *Pseudomonas aeruginosa* were identified in Uganda. Thus, the presence of high abundance of E. coli and ESKAPE pathogens could explain the high number of circulating resistomes in Uganda in contrast to the other East African countries in the study.

In all three countries, Tetracycline antibiotic resistance was observed to be the most prevalent. This may be due to the fact that Tetracycline is the most widely used broad spectrum antibiotic in the East African countries (Kimera *et al*., 2020). According to Sangeda *et al*., 2021, the most common antibacterial agents used in Tanzania are aminoglycosides, beta-lactams, quinolones, tetracycline, trimethoprim and sulfonamides, and various antibacterial cocktails, with Tetracycline class topping the list. These various findings closely align with our study results to a great extent as we also found similar resistances to the drugs described by Sangeda *et al*., 2021, in Tanzania cattle samples. This could be the result of the all-time culprit, that is the overuse or incorrect usage of these antibacterial medications.

Read based methods however, further highlighted that aminoglycosides are the common resistance drug class shared by all the three countries followed by tetracyclines. Elfamycins and MLS (macrolides-lincosamides and streptogramins B) resistance was also identified in Kenya and Tanzania using read based methods. These findings were not featured in assembly-based methods hence highlighting the strength of detecting more AMR genes that read based methods has over assembly-based methods in detection of AMR genes.

The genes identified to be common in the three countries (*aph6-id, tet, sul2, cfxA_gen*) are of public health significance. In several studies, the gene, *aph6-id* which is an aminoglycoside resistance gene has been linked to multi-drug resistant *E. coli* in cattle farms in Korea (Belaynehe *et al*., 2017), in Jeddah (Yasir *et al*., 2020) and in Pakistan (Ali *et al*., 2021). This gives an indication of the widespread distribution of this gene and is a critical epidemiological marker that could increase multidrug resistance in other pathogens through horizontal gene transfer. The genes, *tet and sul2* enhance resistance to sulfonamides and tetracyclines, which are very important antimicrobial classes according to WHO’s ranking (WHO, 2017). These antibiotics are heavily used in livestock industry in East Africa (Orwa *et al*., 2017, Sangeda *et al*., 2021), and over time, the antibiotic residues in beef from cattle may constitute a health risk to humans when consumed. The presence of *cfx* gene, which encodes for cephalosporin resistance indicates that Cephalosporin is also one of the most widely used antibiotics in East Africa. Altogether, this suggests the need for approaches to combat the increasing antimicrobial resistance incidences.

Functional annotation was performed using SqueezeMeta pipeline, which generated heat maps of genes identified in the three EAC countries - Kenya, Uganda and Tanzania. Of the 15 annotated KEGG orthologues, four exhibited significantly elevated gene expression profiles corresponding to the sample genes in the three countries. In Kenya and Tanzania, gene profile K02014 - iron complex outer membrane receptor protein gene, were highly expressed for some samples as seen in Supplementary Figs 3a and 3b. This functional profile gives survival advantage to the bacteria by helping bacteria bind to iron-containing molecules in the environment, thus enabling them to live longer (Palmer and Skaar, 2016 and Caza and Kronstad, 2013), thus potentially leading to the transfer of ARGs or acquiring more ARGs through mutations and horizontal gene transfer.

In Uganda, three KEGG orthologues happened to have a significantly high gene expression profile for some samples as seen in Supplementary Fig 3c. These include; functional profiles K06147 - ATP-Binding cassette subfamily B, K07497 - putative transposase and K07483 - transposase, in their respective decreasing expression levels in TPM (Transcripts per million). According to Vigil-Stenman *et al*., (2017), transposons play a fundamental role in bacterial genome-plasticity and host-adaptation, although their transcriptional activity in the natural bacterial communities has not been largely explored. Thus, the K07497 - putative transposase and K07483-transposase functional profiles as per the KEGG orthologue heat map in Supplementary iFg 3c, suggests the possibility that the introduction and use of antibiotics triggered the changes in genomic regions (mediated by transposons) of the bacteria in the samples. Changes in these regions might have resulted into resistant genes with significant expression in some of the sample genes collected from Uganda.

The use of short reads sequences led to difficulty in assembling some reads into contigs. Lower depth in the samples for Kenya and Tanzania for the analysis might have contributed to less observations of drug resistance classes compared to Ugandan samples. Moreover, there was not enough gut metagenomic samples we could find for all the three countries for our analysis and thus it hampered having solid conclusion about the resistome status in the three countries.

### Conclusions and next steps

Antimicrobial resistance to drugs and other therapeutics is consistently on the rise, as evidenced by the findings of several studies emphasising the numerous routes by which bacteria harbouring AMR genes may be transmitted to humans. Based on the economic value of cattle in Africa and it being a common source of protein, we focused on the East African Cattle to determine which AMR genes would be found through shotgun metagenomic analysis. Our findings indicate that East African cattle’s gut microbiome has a wide range of antibiotic resistance. Thus, the need is urgent for the development of new antibiotic classes to suppress the effects of bacteria harbouring AMR genes. Furthermore, the potential of transmission from cattle to humans poses a larger threat.

This study gives an overview of the current state of antibiotic resistance in East African countries. This includes the identification of a wide range of ARGs, including ARGs linked to multi drug resistance in E. coli such as *aph6-id*. It also associates high prevalence of ARGs to the high abundance of *Enterobacteriaceae* and ESKAPE pathogens. This information could serve as a signal to authorities coordinating East African integration, prompting them to modify and/or formulate policies governing antibiotic use and transit of dairy products across the EAC member nations and Africa at large. It is essential for them to recognize that the development agenda has a cost, which involves focusing and incentivizing researchers to conduct more studies on combating antibiotic resistance, to guarantee a prosperous and harmonious integration of EAC.

Our study acts as a pilot in efforts of AMR surveillance using metagenomics samples in EAC region. Here, we benchmark assembly-based methods against read-based methods. Based on our two workflows, we noted that read based methods is more robust for identification of AMR genes. However, this might vary depending on your research questions and design.

## Availability and Requirements

### Project name

Investigating Antimicrobial Resistance Genes in Kenya, Uganda and Tanzania Cattles Using Metagenomics

### Project home page

https://github.com/omicscodeathon/amr_cattle

### Operating system(s)

Linux CLI

### Programming language

Bash, R 4.2, Perl, Python. **Other requirements:** Java JDK, Chrome, R Studio.

### Licence

MIT

### Any restrictions to use by non-academics

None.

## Availability of data and materials

The dataset analysed during the current study as a case study is publicly available at https://github.com/omicscodeathon/amr_cattle/data. The data supporting the results reported in this manuscript is included within the article and its additional files. The generated progress reports are in HTML format and can be viewed using any preferred browser such as Chrome, Safari, Internet Explorer and/or Firefox. The Project repository which also includes the entire code and other requirements can be downloaded from https://github.com/omicscodeathon/amr_cattle. The guidelines for implementing this tool and related updates, are available at: https://github.com/omicscodeathon/amr_cattle/blob/master/README.md.

## Abbreviations

AMR: Antimicrobial Resistance
ARGs: Antibiotic Resistance Genes
CLI: Command-Line Interface
EAC: East African Community
FDR: False Discovery Rate
EAC: East African Community
GUI: Graphical User Interface
ESKAPE: *Enterococcus faecium, Staphylococcus aureus, Klebsiella pneumoniae, Acinetobacter baumannii, Pseudomonas aeruginosa*, and *Enterobacter spp*

## Acknowledgements

The authors thank the National Institutes of Health (NIH) Office of Data Science Strategy (ODSS), the National Center for Biotechnology Information (NCBI) and the Genetics Society of America for their immense support before and during the October 2022 Omics codeathon organised in collaboration with the African Society for Bioinformatics and Computational Biology (ASBCB).

## Authors Information

**Affiliations**

**Pwani University, Kilifi, Kenya**

Kauthar M. Omar, George L. Kitundu and Felix Lisso

**University of the Western Cape, Cape Town, South Africa**

Abiola A. Babajide

**Institute of Infectious Disease and Molecular Medicine, University of Cape Town, South Africa and National Biotechnology Development Agency, Abuja, Nigeria**

Adijat O. Jimoh

**International Centre of Insect Physiology and Ecology, Nairobi, Kenya**

Dorcus N. Namikelwa

**Ladoke Akintola University of Technology Ogbomoso, Nigeria**

Seun E. Olufemi

**African Society for Bioinformatics and Computational Biology, South Africa Department of Computer Science, University of Ibadan, Ibadan, Oyo State, Nigeria**

Olaitan I. Awe

## Authors Contributions

KO conceived the original idea. KO, GK and FL developed the metagenomics pipeline for Investigating Antimicrobial Resistance Genes in Kenya, Uganda and Tanzania Cattles, performed the bioinformatic analysis of the case study data, and drafted the manuscript. AJ, SO, AB and DN contributed in writing, reviewing and editing the manuscript. OIA’s role was in the administration and supervision of the bioinformatics analysis in the project. OIA also provided the resources to facilitate and complete the analysis and provided guidance. OIA did the editing and final review of the manuscript and provided critical feedback that helped shape the final version of the manuscript. All authors read and approved the final manuscript.

## Declarations

### Ethics approval and consent to participate

Not applicable.

### Consent for publication

Not applicable.

