## Supplementary figures and images for "Investigating Antimicrobial Resistance Genes in Kenya, Uganda and Tanzania Cattle Using Metagenomics"

### Supplementary Figure 1

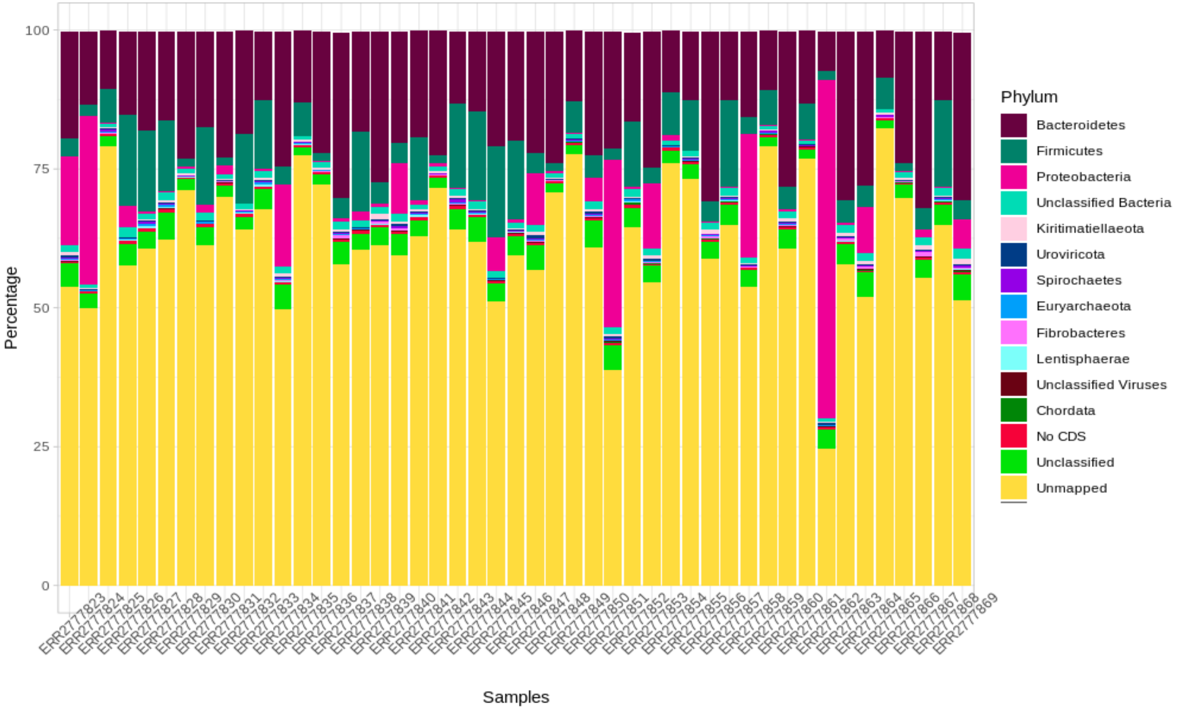

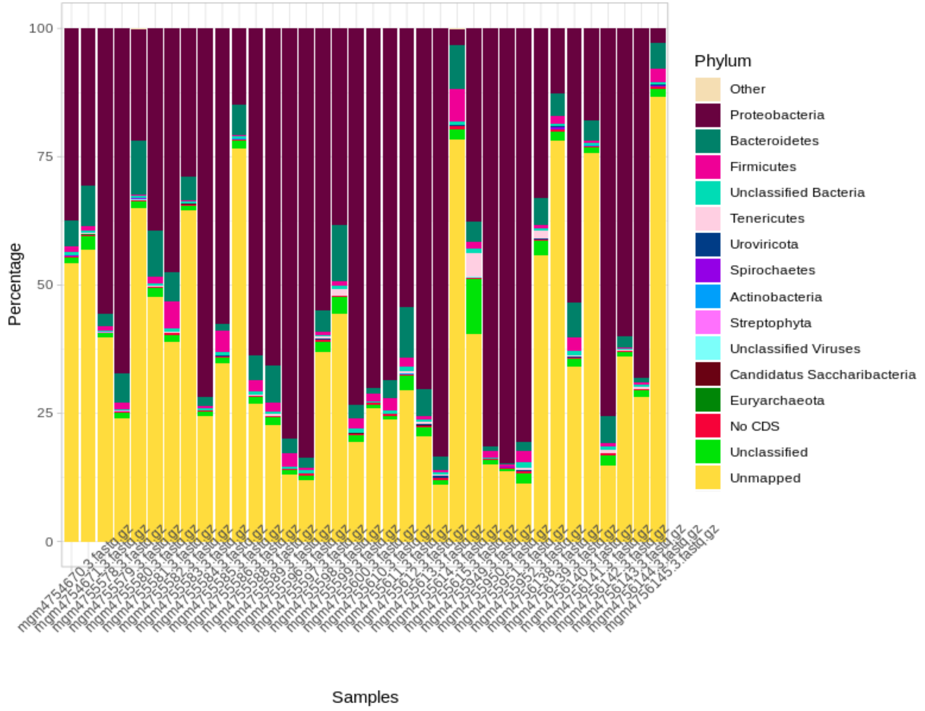

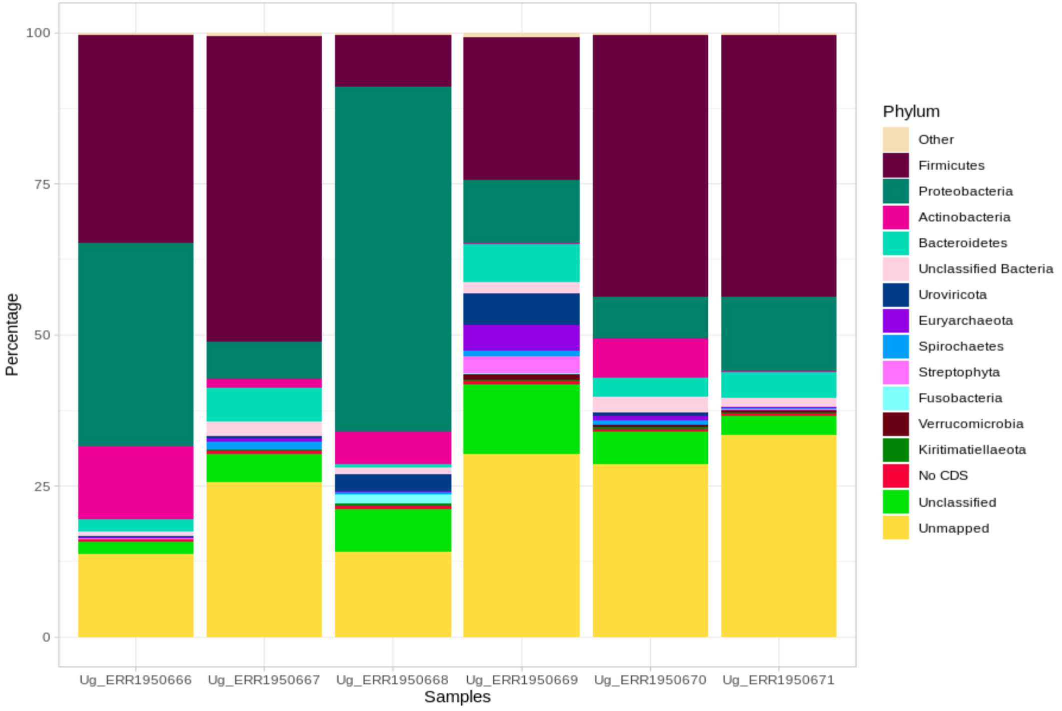

### Supplementary figure 2

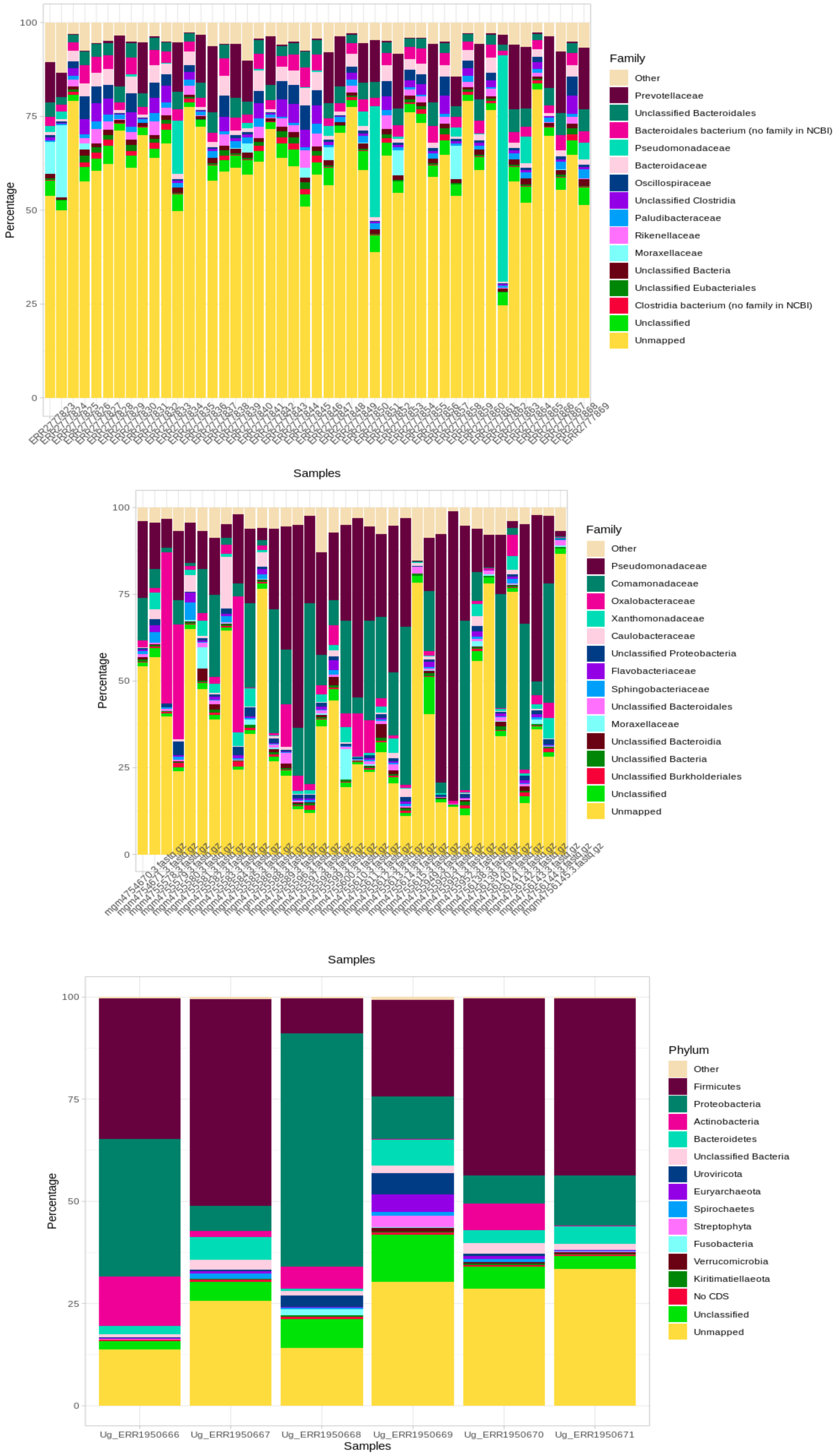
