## Supplementary material for "Investigating Antimicrobial Resistance Genes in Kenya, Uganda and Tanzania Cattle Using Metagenomics"

**Supplementary methods: Data Search**

The data used was searched using advanced search with the following keywords and option: ((((((cattle metagenomics) AND (antimicrobial resistance)) OR (Kenya)) OR (Uganda)) OR (Tanzania)) OR (South Sudan)) OR (Burundi)) OR (Rwanda). A total of 89,515 results were obtained. However, the articles did not contain cattle metagenomic data with links to public databases in which they could be retrieved. I therefore resorted to using google search engine and systematically checking for a paper from East Africa on cattle metagenomics and an aspect of antimicrobial resistance. Several hits were found however the papers with links to public datasets were those for PRJEB28482 (doi: 10.5897/IJLP2019.0654) and PRJEB20456 (doi: 10.1111/tbed.13024) from NCBI and project mgp81260 (<https://www.mg-rast.org/linkin.cgi?project=mgp81260>) from MG-RAST.

**Supplementary methods: Use Cases in the Pipeline**

While using the SqueezeMeta pipeline, we encountered some limitations in the “binning” stage (Step 14). Bins could not be generated from the Kenya and Tanzania dataset while in sequential mode, and so we resorted to using the “co-assembly” mode for the two countries. This means that we treated Kenya and Tanzania as individual sample blocks. Thus, we inferred that this binning limitation was due to the low number of reads in many of the samples from the two countries, given that most of the technical approaches implemented to resolve it failed. However, it is worth noting that we were unsure if that was the actual cause of the Binning failed step.

In analysis using YAMP, we encountered an issue when performing the step, “profile_function”, and this was linked to incompatibility of the downloaded UNIREF database (using YAMP github wiki instructions) with the DIAMOND version in the YAMP pipeline docker image. Retrieving the database from the YAMP MetaPhIAn docker image also proved challenging technical-wise. To address this, we built the compatible database from scratch using a script provided in our github repository (script name: scripts/YAMP/build_diamond.sh). The results obtained from YAMP were similar to those obtained from the SqueezeMeta pipeline. Additionally, YAMP resolved some unmapped regions seen in SqueezeMeta. YAMP results were obtained in tsv files and visualisation of this output was challenging as it was not compatible with most visualisation software and tools. We therefore opted to use a custom pipeline for taxonomical assignment using centrifuge for the analysis and krona for visualisation. The custom pipeline was faster than SqueezeMeta and YAMP consuming less computational resources than both. Further instructions on these use cases are found in our github repository.

The AMRplusplus tool for mining ARGs from reads was developed for paired end reads. However, in our dataset, this comprised of single end reads from Kenya and Tanzania. We therefore customised the AMRplusplus source code to run on our single end reads. We uploaded the single end AMRplusplus scripts on our github repository for reproducibility of the analysis when running on single end reads.

**Supplementary Fig1: Taxonomic abundance at Phylum level for the assembled contigs.** Taxonomic assignment of the contigs was done by Squeezemeta using RDP classifier. The contigs were obtained from Squeezemeta using MEGAHIT assembler. Significant number of contigs did not map to the classifier database (unmapped). (a) Kenyan samples were co-assembled. *Bacteroidetes, Firmicutes* and *Proteobacteria* appear to be the most abundant phyla (b) Tanzania sampless were co-assembled. *Proteobacteria* and *Bacteroidetes* appear to be the most abundant phyla (c) Uganda samples were sequentially assembled. *Firmicutes* and *Proteobacteria* appear to be the most abundant phyla

**Supplementary Fig2: Taxonomic abundance plot at family level for the assembled contigs.** Taxonomic assignment of the contigs was done by Squeezemeta using RDP classifier. These are the contigs obtained from Squeezemeta using MEGAHIT assembler. Significant number of contigs did not map to the classifier database (unmapped). (a) Kenyan samples were co-assembled. (b) Tanzania samples were co-assembled. (c) Uganda samples were sequentially assembled.

**Supplementary Fig 3:** **Heat Map of KEGG Orthologues describing the functional annotation of samples.** The heatmap was as produced by SqueezeMeta. The shading of the heat map is such that, the darker the shade of a pathway, the more abundant that pathway is in that particular sample. (a) Kenyan samples. (b) Tanzanian samples. (c) Ugandan samples.

**Supplementary Fig 4: A Pie Chart describing drug classes composition for AMR Genes mined from Assembled Contigs.** The pie chart was generated in R from the output of Abricate selecting only the drug class column (a) Kenyan samples. (b) Tanzanian samples. (c) Ugandan samples.

**Supplementary Fig 5: Venn diagram showing the total number of drug classes identified per country and in comparison, across the three East African countries from AMR genes mined from reads.** The Venn diagram was created using the bioinformatics and evolutionary genomics Venn diagram tool from a text file containing AMRplusplus output. The output files were filtered using the Linux command cut to only include the drug class column. A total of 59 distinct drug classes were found in the three countries, including all identified 59 in Uganda, 44 out of 56 in each country in Kenya and Tanzania.

**Supplementary Table 1: Drug classes in which resistance was identified in Kenya, Uganda and Tanzania from AMR genes mined from reads.** The table was created using the bioinformatics and evolutionary genomics Venn diagram tool beneath text findings from an AMRplusplus output text file. The output of AMRplusplus was filtered using the Linux command cut to only include the column for drug class. This table supplements Supplementary Fig 5 by giving names to the number of drug classes shown in the Venn diagram

**Supplementary Fig 6: A venn diagram describing the total number of AMR Genes in Kenya, Uganda and Tanzania uniquely as well as comparisons of genes available across the three countries mined from reads.** The Venn diagram was generated from an AMRplusplus output text file using the bioinformatics and evolutionary genomics Venn diagram tool. The result files were filtered using the Linux command cut to only include the column for AMR genes. across total, 1420 unique AMR genes were found across the three countries, with 965 in Uganda, 403 in Tanzania, and 353 in Kenya.

**Supplementary Table 2: AMR Genes present in Kenya, Uganda and Tanzania.** Counts in Individual Countries Indicate the Genes that are Unique to that Country mined from reads. The table was built using the Venn diagram tool for bioinformatics and evolutionary genomics under text findings from an AMRplusplus output text file. The AMRplusplus output was filtered using the Linux command cut to contain only the column for AMR genes. This table supplements Supplementary Fig 6 by naming the AMR genes that occur in the Venn diagram.
